# Cell Type-Agnostic Optical Perturbation Screening Using Nuclear In-Situ Sequencing (NIS-Seq)

**DOI:** 10.1101/2024.01.18.576210

**Authors:** Caroline I. Fandrey, Peter Konopka, Marius Jentzsch, Afraa Zackria, Salie Maasewerd, Eicke Latz, Jonathan L. Schmid-Burgk

## Abstract

Genome-scale perturbation screening is widely used to identify disease-relevant cellular proteins serving as potential drug targets. However, most biological processes are not compatible with commonly employed perturbation screening methods, which rely on FACS- or growth-based enrichment of cells. Optical pooled screening instead uses fluorescence microscopy to determine the phenotype in single cells, and subsequently to identify individual perturbagens in the same cells. Published methods rely on cytosolic detection of endogenously expressed barcoded transcripts, which limits application to large, transcriptionally active cell types, and often relies on local clusters of clonal cells for unequivocal barcode assignment, thus precluding genome-scale screening for many biological processes. Nuclear In-Situ Sequencing (NIS-Seq) solves these shortcomings by creating bright sequencing signals directly from nuclear genomic DNA, enabling screening any nucleus-containing cell type at high density and high library complexity. We benchmark NIS-Seq by performing three genome-scale optical screens in live cells, identifying key players of inflammation-related cellular pathways.

## INTRODUCTION

Genetic perturbation screens aim to decipher the relationship between genotype and phenotype by ablating random genes in a pool of cells. Perturbation screens are commonly based on enrichment of cells with a phenotype of interest followed by next generation sequencing of the perturbagens, e.g., CRISPR sgRNAs ^1 2^ or gene trap insertions in haploid cells ^3^. Enrichment of specific perturbagens allows to conclude to the biological function of targeted genes. While enrichment is commonly achieved through cell-growth or fluorescence activated cell sorting (FACS) ^4 5^, more sophisticated screening approaches were realized combining perturbation screens with single cell molecular profiling via mass cytometry or RNA sequencing ^6 7 8^. Those methods acquire high-dimensional information on single cell level, but are incapable of monitoring dynamic processes due to their disruptive nature.

For arrayed CRISPR screening, perturbed cells are physically separated into compartments containing cells with largely homogeneous genotypes. This allows to analyze cells using complex assays, such as fluorescence microscopy ^9^, and thus to monitor cell physiology in a spatially and temporally resolved manner. However, as arrayed screens are laborious and prone to experimental inconsistencies between compartments, they are mostly feasible for small targeted perturbagen libraries.

Optical pooled perturbation screening addresses these limitations by accessing the bandwidth of phenotypes that can be observed by fluorescence microscopy while working with a complex pool of cells ^10^. Currently, several approaches to optical pooled perturbation screening were developed: For Optical-Based Enrichment, cells with a phenotype of interest are identified by microscopy and are marked by light-based conversion of a photoactivatable fluorescent protein. Subsequently, cells are dissociated, marked cells are enriched by FACS sorting, and perturbagens are analyzed by deep sequencing ^11^. To a similar end, image-based cell sorting ^12^ or laser microdissection ^13^ can be used to isolate cells displaying a microscopic phenotype of interest for subsequent perturbagen identification, albeit at limited optical resolution. Instead of enriching for a population of cells for lysis and sequencing, Feldman et al. ^10^ developed a method for microscopic perturbagen identification based on in-situ sequencing of barcodes contained in cellular mRNAs, using signal amplification by rolling circle amplification (RCA). Using this method, both the phenotype and the genotype are recorded from individual cells using fluorescence microscopy, allowing to categorize cellular phenotypes post-hoc ^14^. This method was recently adapted to screen genome-scale perturbation libraries for pre-defined ^15^ and multi-dimensional phenotypes ^16 17^.

While in-situ sequencing-based screening has the advantage of post-hoc genetic dissection of multi-dimensional phenotypes, current methods still bear two major limitations: Because sequencing is dependent on mRNA molecules in the cytosol, cells with low transcriptional activity or small volume generate insufficient signal, precluding many relevant cell types from optical pooled screening. Furthermore, cells have to be permeabilized for in-situ sequencing. Thus, loss of precise cell-cell boundaries can interfere with assignment of cytosolic spots to nuclei at high cell densities and complex barcode distributions.

To address these shortcomings, we developed nuclear in-situ sequencing (NIS-Seq): After phenotyping live cells and fixation, the reverse-complement sequences of perturbation gRNAs are locally transcribed from genomic DNA using T7 polymerase by an adapted *Zombie* protocol ^18^. Nuclear clusters of RNA are efficiently sequenced by padlock-based in-situ sequencing. NIS-seq circumvents all major drawbacks of pooled optical screening by unambiguously assigning bright signal clusters to nuclei independent of cell size, type, or transcriptional activity.

## RESULTS

### NIS-Seq enables cell type-agnostic optical barcode identification

Based on adapting two previously published methods ^10 18^, we developed nuclear in-situ sequencing (NIS-Seq) to enable cell-type-agnostic high-density optical CRISPR screening. While published in-situ sequencing detects barcodes from RNA Polymerase II-expressed mRNAs in the cytosol, NIS-Seq uses T7 in vitro transcription to generate multiple RNA copies in the nucleus (Figure 1A). After subsequent reverse transcription, padlock elongation, ligation, and rolling circle amplification, sgRNAs are identified by 14 cycles of sequencing-by-synthesis using three excitation wavelengths (Figure 1A). Nuclear signals enable unambiguous assignment to cells even at high cell densities as required for genome-scale screening. Not relying on transcriptional activity or cytosolic volume, NIS-Seq was found compatible with THP1-derived and primary human macrophages, whereas the previously published in-situ sequencing protocol failed in these cell types (Figure 1B). For cell-accurate mapping of live cell phenotyping data to NIS-Seq data using different objectives and high cell densities, we optimized a cross correlation-based search algorithm that can compensate small movements and distortions of cells and glass plates during live-cell imaging, assigning pairs of nuclei between imaging modalities with high confidence (Figure 1C). We next generated a genome-scale perturbation library in THP1-derived macrophages. Sequencing this library using NIS-Seq revealed clean nuclear signal intensities over the course of 14 cycles (Figure 1D), which can be translated to known sequences of library members (Figure 1E). Overall, more than 60% of nuclei were unambiguously assigned to single library members with >66% of aggregated spot intensities (Figure 1F, see Methods section for details), while <20% of nuclei had no spots mapping to the library. Overall, library representation was highly correlated between NGS- and NIS-Seq based sequencing (Fig. 1G). Finally, to ensure high genome efficiencies using a NIS-Seq-compatible lentiviral CRISPR vector, we compared different vector designs across three genomic target sites in HeLa-Cas9 cells. Genome editing efficiencies were determined by NGS after Puromycin selection and Cas9 induction ^19^, confirming that inserting an inverted T7 promoter after the Polymerase-III terminator of a sgRNA expression cassette mirrors efficiencies of state-of-the-art lentiviral CRISPR vectors (Figure 1H).

**Figure 1.**
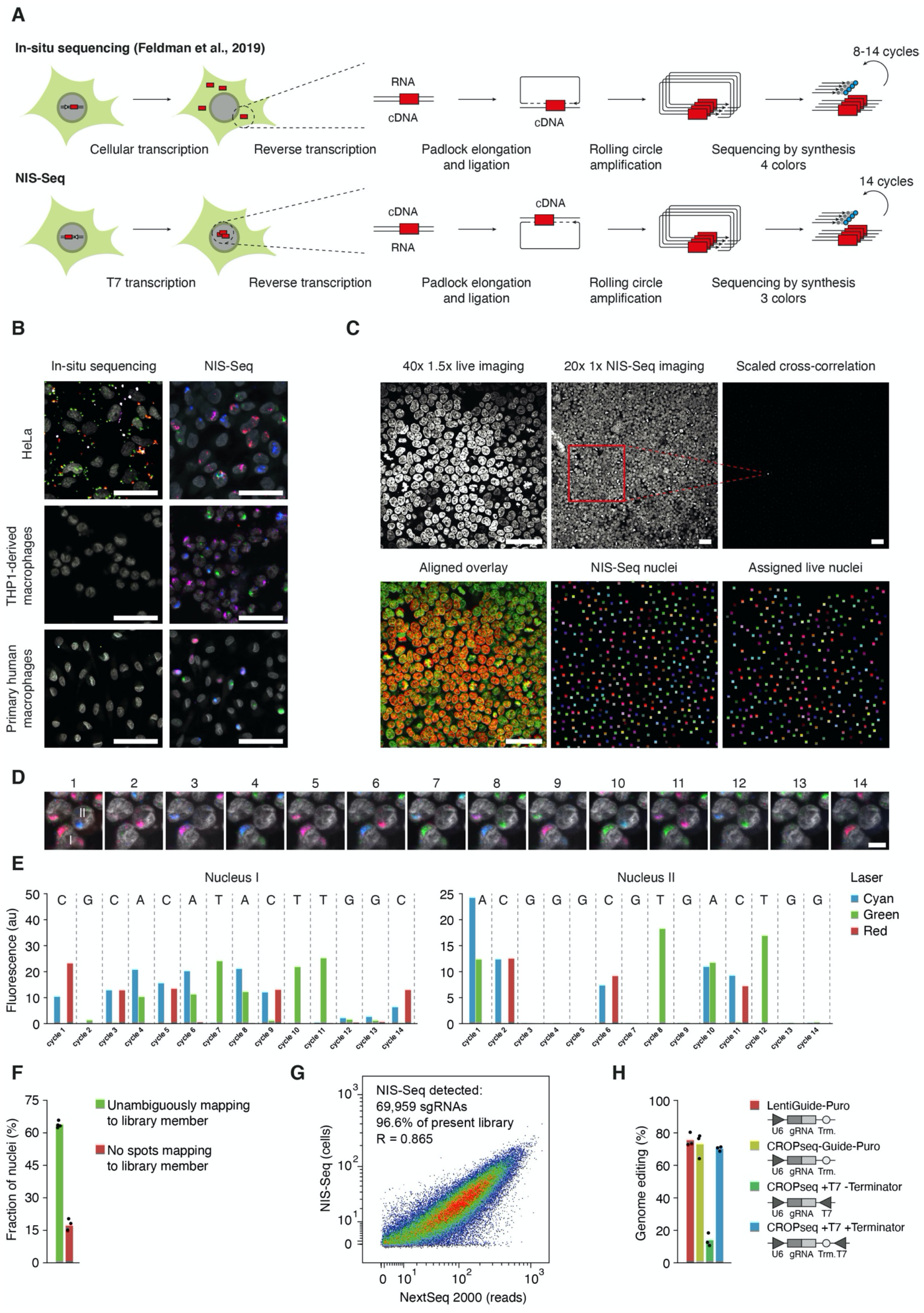
NIS-Seq enables optical barcode identification in nucleated cells. **(A)** Outline of NIS-Seq reaction steps in comparison to previously established in-situ sequencing of barcoded mRNA ^10^. **(B)** NIS-Seq imaging results in comparison to cytosolic in-situ sequencing results obtained across three cell types. Scale bar, 50 μm. **(C)** Nuclei assignment between live cell imaging and NIS-Seq data across different objectives and timepoints. Nuclear staining patterns are mapped by two-dimensional FFT-accelerated high-pass filtered cross-correlation (top right panel). Overlay of nuclear signal reveals slight dislocation of nuclei between imaging timepoints (bottom left). Centers of gravity of CellPose-defined nuclei are assigned to nearest neighbors and ambiguous assignments are removed (bottom center and right). Assigned nuclei are color-coded by the same random color. Scale bar, 50 μm. **(D)** Raw images of 14 cycles of NIS-Seq barcode sequencing. Nuclear staining was performed at cycles 1, 4, 7, 10, and 13. Scale bar, 10 μm. **(E)** Quantitative spot intensities obtained from nuclei I and II highlighted in (D). Indicated on top is the base calling result, matching two members of the pooled lentiviral library used. **(F)** Fraction of nuclei mapping to known library member sequences. Unambiguous mapping is defined as more than two thirds of aggregated library-matching spot intensities within a nucleus mapping to a single library member. **(G)** Library coverage in transduced THP1 macrophages, measured by PCR-based NGS and NIS-Seq. Each library member covered in PCR-based sequencing is represented by one dot; dropping out sgRNAs, e.g., those targeting essential genes, are not shown. Jitter is added to values to visualize spot density even at low integer values. **(H)** Genome editing efficiencies of widely-used sgRNA-expressing lentivirus designs compared to constructs with an additional T7 promoter inserted for NIS-Seq in reverse-orientation. Genome editing efficiencies at three independent loci were assessed by NGS ^19^ in HeLa-Cas9 cells transduced with indicated lentiviral constructs after four days of Puromycin selection and Cas9 induction.

### Genome-scale screening in HeLa cells reveals known components of two immune receptor signaling pathways

To demonstrate genome-scale screening applications of NIS-Seq, we targeted HeLa cells to investigate genes involved in the activation of the Nuclear Factor (NF)-κB, a family of five members of inducible transcription factors functioning as homo- or hetero-dimers ^20^. In an inactive state, NF-κB dimers are retained in the cytosol by inhibitory proteins such as IκB. Upon activation of the pathway by external stimuli, phosphorylation of the Iκk complex leads to ubiquitination and proteasomal degradation of IκB, allowing dimeric NF-κB to shuttle into the nucleus ^20^. To study genes involved in this process, we used a previously established translocation assay based on HeLa cells carrying a fluorescent p65-mNeonGreen reporter ^10^. To elucidate pathway members, we performed two replicate screens using 76,441 sgRNAs targeting human protein coding genes as well as 1,000 non-targeting control guides. HeLa cells were stimulated with interleukin 1 β (IL-1 β) or tumor necrosis factor alpha (TNF-α) to target two different pathways of NF-κB activation. After live cell phenotyping, the sgRNA identity present in each cell was determined by NIS-Seq. Mapping of phenotype and NIS-Seq nuclei resulted in >14,000 genes covered by at least 15 cells in each of two replicate screens for each stimulus (Figure 2A, D). Nuclear translocation of p65 was quantified as pixel-wise Pearson correlation coefficient between p65-mNeonGreen and nuclear staining images. Genes with altered mean nuclear translocation across targeted cells corresponded to the receptors IL1R1 and TNFRSF1A as well as multiple known downstream pathway members such as TRAF6, CHUK, and IKBKG, which were confirmed by single-cell collages of targeted cells retrieved from pooled imaging data (Figure 2B, E). Strong outlier genes were furthermore confirmed using orthogonal sgRNAs from the Toronto KO CRISPR library v3 ^21^ (Fig. 2C, F).

**Figure 2.**
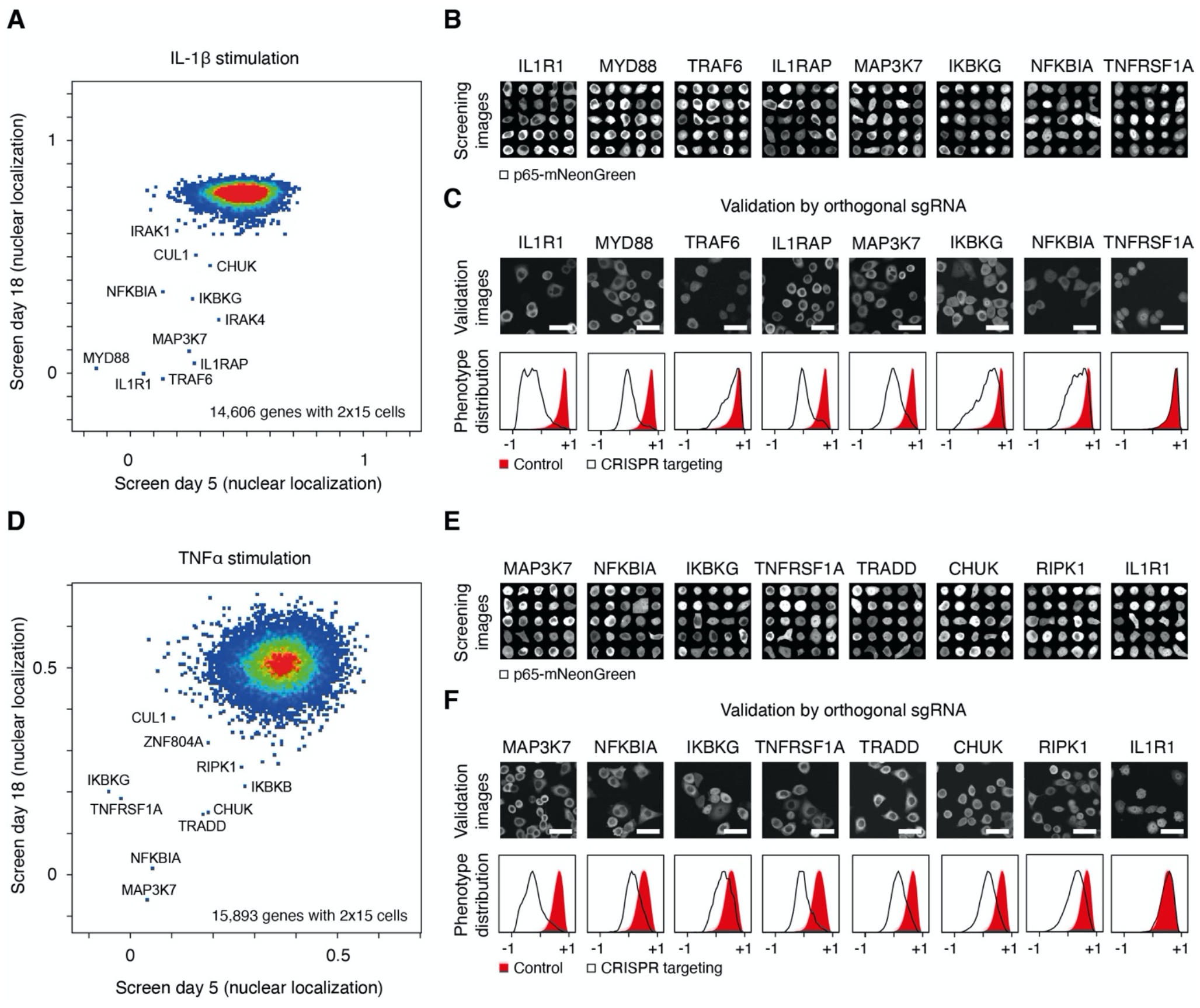
Genome-scale optical perturbation screening for mediators of NF-κB activation. **(A)** Results of two replicates of genome-scale NIS-Seq perturbation screening in HeLa-Cas9-p65-mNeonGreen cells stimulated with IL-1β. Dots correspond to genes and axes indicate mean pixel-wise Pearson correlation between mNeonGreen and nuclear staining signals. **(B)** Collages of cellular images from (A) mapped to perturbed genes indicated. Shown is the mNeonGreen signal.**(C)**Arrayed hit validation in HeLa-Cas9-p65-mNeonGreen cells using alternative sgRNA sequences from the Toronto KO library v3. Top panel, exemplary mNeonGreen images of IL-1β-stimulated cells. Bottom panel, distribution of activation states, quantified by pixel-wise Pearson correlation between mNeonGreen and nuclear staining signals. Scale bar, 50 μm. **(D)** Results of two replicates of genome-scale NIS-Seq perturbation screening in HeLa-Cas9-p65-mNeonGreen cells stimulated with TNF-α. Dots correspond to genes, and axis positions indicate mean pixel-wise Pearson correlation between mNeonGreen and nuclear staining signals. **(E)** Collages of cellular images from (D) mapped to perturbed genes indicated. Shown is the mNeonGreen signal. **(F)** Arrayed hit validation in HeLa-Cas9-p65-mNeonGreen cells using alternative sgRNA sequences from the Toronto KO library v3. Top panel, exemplary mNeonGreen images of TNF-α-stimulated cells. Bottom panel, distribution of activation states, quantified by pixel-wise Pearson correlation between mNeonGreen and nuclear staining signals. Scale bar, 50 μm.

### Genome-scale optical screening for mediators of inflammasome activation in human monocytes

NLRP3 inflammasomes are megadalton protein complexes that assemble in response to danger- and pathogen-associated molecular patterns in macrophages, leading to rapid IL-1 β and IL-18 cytokine release as well as a rapid form of programmed cell death termed pyroptosis. Even though this pathway is critically involved in a multitude of age-related diseases, no genetic screening has been performed in human cells to systematically identify genetic components involved in NLRP3 inflammasome assembly. Using NIS-Seq, we screened a genome-scale CRISPR KO library of THP1 monocyte-derived macrophages stimulated with the ionophore Nigericin. The cells used were deficient in Caspase 1 and 8 in order to avoid pyroptotic cell death downstream of early steps of inflammasome assembly. An ASC-GFP reporter enabled monitoring inflammasome assembly in live cells. Inflammasome activation was quantified in individual cells by acquiring Z-stacks, and calculating the ratio of overall GFP intensity to high-pass filtered GFP intensity, the latter originating from smaller objects like ASC specks. Correlating the results of two independent screens in live macrophages, we confirmed NLRP3 and IKBKB as the most critical pathway members ^22^, while ablation of MAP3K7, TRAF6, HSP90B1, or RNF31 resulted in intermediate degrees of pathway dysfunction (Fig. 3A). Aggregated images of live cells assigned to hit genes confirmed a reduction in ASC specking as compared to control cells (Fig. 3B). While the relevance of the HSP90 chaperone for inflammasome activation had been observed before ^23 24^, a specific involvement of the beta isoform has not been described, and was not observed in previous genetic screens in murine macrophages ^25^. Independent validation of hit genes by lentiviral expression of orthogonal guide RNAs from the Toronto Knock Out (TKO) library confirmed reduced ASC specking in response to Nigericin (Fig. 3C), which was reflected in cytokine secretion levels in a clonal NLRP3 knock-out cell line (Fig. 3D).

**Figure 3.**
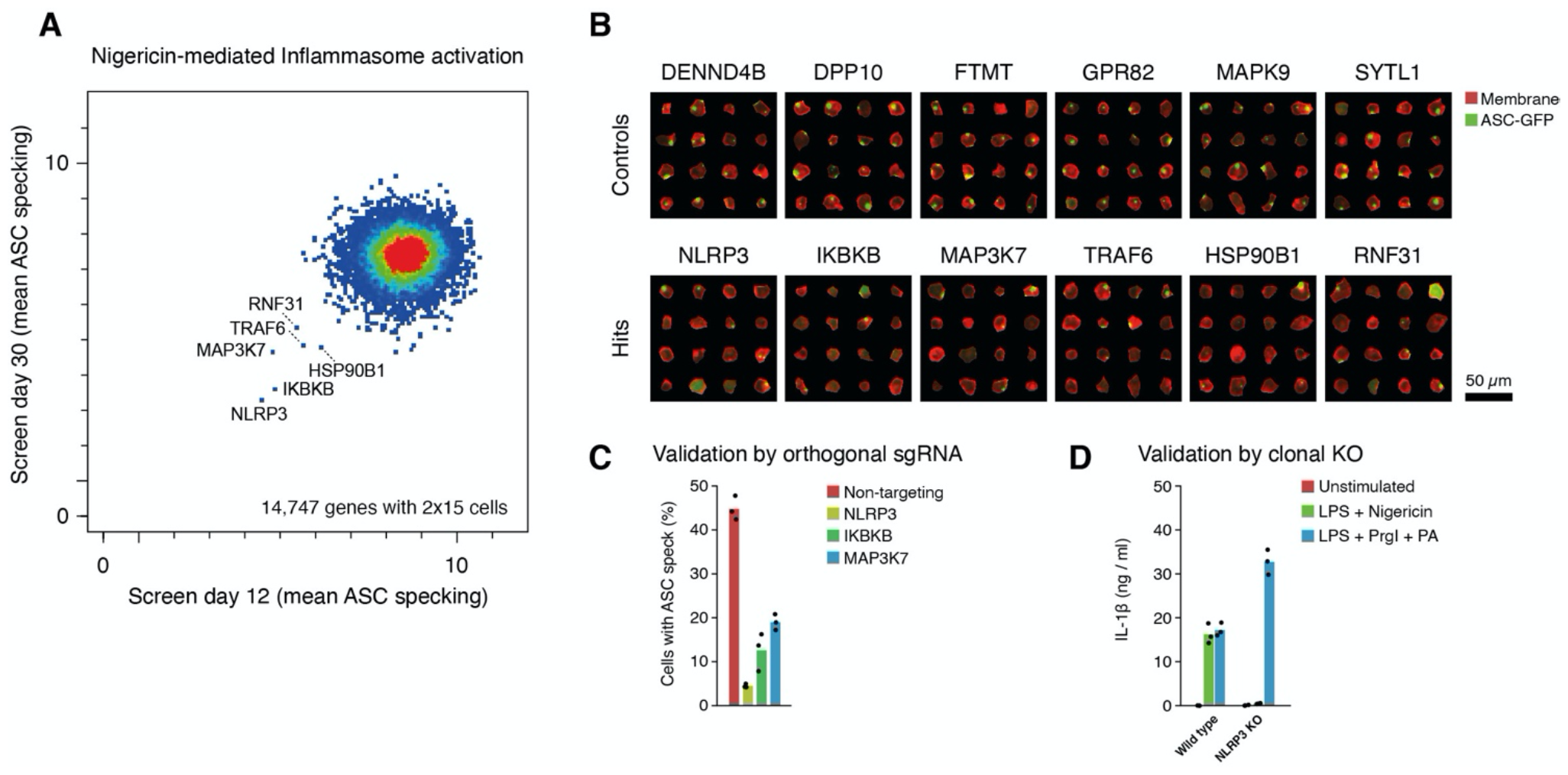
Genome-scale optical perturbation screening in THP1-derived macrophages for inflammasome activation. **(A)** Results of two replicates of genome-scale NIS-Seq perturbation screening in THP1-Cas9-ASC-GFP-CASP1/8^DKO^ cells stimulated with Nigericin. Dots correspond to genes and axes indicate mean ratios of high-pass filtered GFP signal relative to the overall GFP signal per cell. **(B)** Collages of cellular images from (A) mapped to perturbed genes indicated. Shown are membrane stain (red) and ASC-GFP (green) signals. **(C)** Arrayed hit validation in THP1-Cas9-ASC-GFP cells using alternative sgRNA sequences from the Toronto KO library v3. Shown are fractions of cells with an ASC speck upon Nigericin stimulation in three replicate wells. **(D)** Cytokine secretion in response to two inflammasome triggers in wild type or clonal NLRP3-deficient THP1-ASC-GFP cells, measured by IL-1β ELISA. Shown are three replicate wells of stimulated cells.

## DISCUSSION

While efficient editing of single genomic loci for functional genomics studies was possible with predecessor technologies of CRISPR, genome-scale perturbation screening has been revolutionized by the programmability of CRISPR nucleases through short lentiviral sgRNA expression cassettes. Perturbation screening enables to systematically discover genetic determinants of biological processes ^4 5 6 7 8^. NIS-Seq significantly extends the applications of perturbation screening by enabling optical phenotyping in living cells of any nucleated type. It is compatible with high cell density, high library complexity and highly dynamic phenotypes. Thus, biological processes can be mapped to involved genes quantitatively and kinetically, not only identifying critical pathway members, but also reading out genetic rheostats or pathway intersections. NIS-Seq is expected to be compatible with any phenotype observable by microscopy, including sub-cellular transport, cell migration, dynamic oscillation, protein complex formation, RNA splicing, or single-molecule RNA localization ^26^, optionally using newly developed super-resolution ^27^ or non-optical microscopy techniques ^28^. Ongoing improvements in image acquisition speed will be critical to observe sufficient numbers of cells and to enable reliable assignment of nuclei between phenotyping and sequencing images. To allow for longer live imaging times, an imaging sequence could be deployed acquiring low-magnification overview images in regular intervals between high-resolution imaging, keeping track of cellular movements. NIS-Seq will not only enable loss-of-function screening, but also enable CRISPR-activation or cDNA overexpression profiling, which could help to identify optimized iPS-cell differentiation protocols based on optical cell type identification ^29^.

NIS-Seq requires five days of hands-on time for performing a genome-scale screen. Many manual steps currently rely on specific pipetting techniques and are very repetitive; to enable future scale-up, faster cycling times, and to minimize reagent use, we have successfully automated all repetitive steps using an off-the-shelf pipetting robot. To reduce acquisition times of NIS-Seq sequencing cycles, we successfully tested base calling with laser-only switching using a multiband emission filter. In the future, antibody-based base detection might further increase signal intensity, reduce amplification steps, or reduce phasing artifacts ^30^.

All NIS-Seq image analysis can performed using our open-source online analysis applications (*jsb-lab*.*bio/opticalscreening*), which does not require local software installation, server hardware or coding experience to analyze genome-scale optical perturbation screens.

## Acknowledgements

We thank David Feldman and Thomas Ebert for helpful discussions and Lukas Rossnagel, Marta Lovotti, Bernardo Franklin, Florian Schmidt, and Florian Gohr for help with primary human monocytes. The HeLa-Cas9-p65-mNeonGreen cells were kindly provided by the Blainey Lab and the Cheeseman Lab ^31^. J.S.-B. was supported by Deutsche Forschungsgemeinschaft (DFG) under Germany ‘s Excellence Strategy – EXC2151 – 390873048, Deutsche Forschungsgemeinschaft (DFG) SFB 1454 - project 432325352, and TRA Life & Health (University of Bonn).

## Author contributions

J.L.S.-B. conceived of the study. C.I.F., P.K., M.J., and J.L.S.-B. designed the experiments. C.I.F., M.J., P.K.,A.Z., and S.M. collected the data. C.I.F., P.K., and J.L.S.-B. analyzed the data. C.I.F., P.K., E.L., and J.L.S.-B. wrote the manuscript with input from all coauthors.

## METHODS

### CROPseq-iT7 backbone

CROPseq-Guide-Puro ^8^, which was a gift from Christoph Bock (Addgene #86708), was PCR-amplified with primers CROPseq_iT7_fwd/rev using Q5 polymerase. PCR products were DpnI-digested, 5 ‘-phosphorylated, and circularized by T4 ligation. Transformants were sequence-verified using Tn5-mediated whole-plasmid tagmentation and MiSeq sequencing. The NIS-Seq-compatible CROPseq-iT7 backbone was deposited to Addgene (#211699).

### Library cloning

Library cloning was performed as previously described by Joung *et al*. ^32^. In brief, the Human Brunello CRISPR knockout pooled library, a gift from David Root and John Doench (Addgene #73178) ^33^, was PCR-amplified with gRNA_library_fwd/rev primers. The target vector CROPseq-iT7 was digested with the restriction enzyme Esp3I. Subsequently, the purified PCR product was cloned into the digested vector using Gibson assembly. The assembled reactions were pooled and purified via a Zymo DNA Clean & Concetrator-25 column. The purified library was electroporated in eight replicates into Endura electrocompetent cells (Lucigen), using 50-100 ng/μl DNA and 25 μl cells in 0.1 cm BioRad cuvettes at 1800 V, 10 μF and 600 Ω. Cells were directly recovered in 975 μl pre-warmed recovery medium and incubated for one hour at 37°C and 300 rpm shaking. Then, each culture was transferred to 1 l LB medium containing 100 μg/ml ampicillin and grown overnight at 37°C 230 rpm shaking. DNA was purified by maxiprep using a Purelink™ HiPure Plasmid Maxiprep Kit. Library coverage was determined by Illumina Next Generation Sequencing using staggered guide-specific primers for a first target amplification PCR and dual indexing barcodes for a second barcoding PCR using NEBNext PCR polymerase.

### Tissue culture

HeLa cells were cultivated in DMEM Glutamax media supplemented with 10% FCS and 10 μg/ml Ciprofloxacin in a 37°C incubator with 5% CO_2_. THP1 cells were cultivated in RPMI Glutamax media supplemented with 10% FCS and 10 μg/ml Ciprofloxacin in a 37°C incubator with 5% CO_2_. Primary human monocytes were obtained from fresh human blood (obtained as approved by the Ethics Committee of the University Hospital Bonn) by Ficoll gradient centrifugation and subsequent CD14-based MACS separation (Miltenyi, 130-050-201) and cultivated for differentiation in RPMI Glutamax media supplemented with 10% FCS, 10 μg/ml Ciprofloxacin (Sigma), and 2.5 μg/ml M-CSF (ImmunoTools). THP1 cells expressing ASC-GFP from an NF-κB-dependent promoter were purchased from Invivogen (thp-ascgfp).

### Library transduction and quality control

1.5 x 10^7^ HEK 293T cells were transfected in a 15 cm tissue culture dish using Lipofectamine 2000 (Invitrogen) with 14.1 μg Brunello_iT7 plasmid library, 7.0 μg lentiviral packaging plasmid pMD2.G and 10.6 μg lentiviral packaging plasmid psPAX2. After 4-6 h of incubation, the medium was changed. 48 hours later, virus-containing supernatant was filtered through a 0.45 μm filter (Merck Millipore), aliquoted, and stored at - 80°C. For each of four screening replicates, 8 x 10^6^ HeLa-Cas9-p65-mNeonGreen cells ^10^ were transduced with 1 ml lentivirus library and 10 μg/ml polybrene (Merck Millipore). After one day, cells were selected and induced with 3 μg/ml Puromycin and 1 μg/ml Doxycycline (Cayman Chemicals). Cells were split 1:3 when reaching confluency. For each of two macrophage screening replicates, 1.6×10^7^ THP1-ASC-GFP-Cas9 CASP1/8^DKO^ cells were transduced with 2 ml of lentivirus library and 10 μg/mL polybrene. The next day, cells were selected with 3 μg/ml Puromycin. After 4-7 days of selection and induction, one million cells were lysed in 100 μl of direct lysis buffer at 65°C for 10 minutes, and 95°C for 15 minutes ^19^. Genomically integrated guide sequences were amplified using NEBNext PCR polymerase (NEB) and staggered guide-specific primers (Supplementary Table 1) in two replicates per library. After secondary barcoding PCR, purification, and Nanodrop-based quantification, libraries were sequenced on an Illumina NextSeq 2000 using a P2 100-cycle cassette. Library members were counted using the online application www.jsb-lab.bio/LibCounter.htm.

### NIS-Seq

For genome-scale NIS-Seq perturbation screens, 24-well glass bottom plates (GreinerBio) were coated with 0.1% poly-L-Lysine (w/v in H_2_O; Sigma-Aldrich) for 30 minutes at room temperature and washed three times with PBS. For HeLa screens, cells were seeded at 5 or 18 days of doxycycline induction. Per replicate, 4 x 10^5^ HeLa-Cas9-p65-mNeonGreen Brunello-iT7 library cells were seeded per well and incubated overnight. The next day, cells were stimulated for 45 minutes with 30 ng/ml human recombinant IL-1α (rcyec-hil1b; Invivogen) or 30 ng/ml human recombinant TNF-α (rcyc-htnfa; Invivogen), respectively. Live cell nuclei were stained with 2 μM Hoechst 33342 and membranes were stained with 200 ng/ml CellMask Plasma Membrane Stain Deep Red (Thermo). For THP1 screens, THP1-ASC-GFP-Cas9 CASP1/8^KO^ Brunello-iT7 cells were pre-differentiated overnight with 100 ng/ml PMA (Invivogen). The next day, cells were carefully washed, detached, and 8 x10^5^ library cells were seeded per well in PLL-coated glass bottom plates. The next day, cells were primed with 200 ng/ml LPS (Sigma Aldrich) for 3 hours, pre-incubated with 50 μM Z-VAD (MedChemExpress) for 30 minutes and stimulated with 7.5 μg/ml Nigericin (Cayman Chemicals) for 1 hour. Cell membranes were stained with 200 ng/ml CellMask Plasma Membrane Stain Deep Red (Thermo). All live cell phenotype images were acquired in DMEM Fluorobrite with 10 mM HEPES and 2 μM Hoechst 33342 using a 20x objective for HeLa cells and 10x objective with Z-stacks for THP1 cells. After phenotyping, cells were fixed and permeabilized with a 3:1 mixture of methanol and acetic acid for 20 minutes. The fixation was carefully replaced with 1X PBS to avoid dehydration of the cells. Cells were washed with nuclease free water before adding 1x T7 in-vitro transcription mix (T7 MEGAscript, Thermo) for 3 hours at 37°C. After IVT, cells were fixed with 4% paraformaldehyde in PBS for 20 minutes and washed with PBS-T (PBS + 0.1% Tween-20) two times. Reverse transcription and post-fixation steps were performed as described by Feldman *et al*. ^10^ using oRT_CROPseq_iT7 as reverse transcription primer. After post-fixation, cells were washed three times with PBS-T and incubated with a gap-fill Phusion mix (1x Ampligase Buffer, 0.4 U/mL RNase H, 100 nM padlock probe oPD_CROPseq_iT7, 0.0125 U/mL NEB Phusion polymerase, 0.5 U/mL Ampligase, 0.05 mM dNTPs, 0.05 M KCl and 5% formamide) for 30 minutes at 37°C and 45 min at 45°C. Cells were then washed twice with PBS-T and incubated with a rolling circle amplification mix overnight at 30°C ^10^. Cells were washed twice with PBS-T before hybridization of 1 μM of the in-situ sequencing primer oSBS_CROPseq_iT7 in 2x SSC for 5 minutes at 37°C. Cells were washed with MiSeq buffer PR2 (Illumina), and perturbation barcodes were analyzed by sequencing-by-synthesis using reagents from used Illumina NextSeq 2000 P2 cassettes. For each of 14 cycles, cells were incubated with the nucleotide incorporation mix for 3 minutes at 60°C, followed by three rounds of five washes with PR2, each with 5 minutes incubation at 60°C. The incorporated nucleotides were imaged after addition of 200 ng/ml Hoechst 33342 in PR2. After each imaging cycle, fluorescent nucleotides were cleaved and de-blocked by incubation with NextSeq cleavage mix for 3 minutes at 60°C, three washes with PR2, incubation for 2 min at 60°C, and three additional washes before the next cycle of incorporation.

### Imaging

All images were acquired using a Nikon Ti2 body equipped with a Yokogawa CSU-W1 spinning disc unit connected to Lumencor Celesta multimode lasers with wavelengths of 405 nm (nuclear staining), 477 nm (sequencing channel 1, mNeonGreen, and GFP), 546 nm (sequencing channel 2), and 638 nm (sequencing channel 3 and CellMask deep red). Emission filters used were Chroma ET450/50 (nuclear staining), Chroma ET525/50 (sequencing channel 1, mNeonGreen, GFP), 572/28 BrightLine HC (sequencing channel 2), and 680/42 BrightLine HC (sequencing channel 3, CellMask deep red). Exposure times were 90 ms for all channels except p64-mNeonGreen, which was exposed for 150 ms. Objectives used were a Nikon 10x CFI P-Apo, a Nikon 20x CFI P-Apo, or a Nikon 40x CFI Apo 40x WI with or without a 1.5x tube lens inserted into the light path. A Hamamatsu Orca Flash4.0 LT+ camera was used in electronic shutter mode at full resolution (2048×2048).

### Image analysis of NIS-Seq data

Raw images of up to 14 NIS-Seq cycles were aligned by FFT-accelerated cross-correlation of nuclear staining images. Spots were detected by summing up all sequencing channels across the first three cycles, high-pass filtering, local maximum detection, and brightness thresholding. Spot sequence information was aggregated across 5×5 pixels for every spot after high-pass filtering and eliminating negative values. Channel unmixing was performed by multiplying the channel vector of each cycle with the inverse matrix of average base-wise channel intensities. Non-G bases were called by the maximum of unmixed channels, whereas Gs were called at cycles with all unmixed intensities below 20% of the spot ‘s maximum unmixed intensity across all cycles. Sequences were assigned to the dictionary of known sequences (Brunello sgRNA sequences reverse complemented), allowing zero or one mismatch and no ambiguities. Dictionary-matched and -corrected spot sequences were assigned to nuclei, whose outlines were defined by CellPose using the “nuclei” model ^34^, requiring the dominant sequence to make up more than two thirds of total intensity of library spots in a given nucleus.

### Image analysis of live phenotyping data

Z-stacks were collapsed by averaging where applicable. Cell- and nuclear outlines were defined by CellPose using the “cyto2” and “nuclei” models ^34^. Nuclear translocation of mNeonGreen was quantified by calculating the Pearson correlation between nuclear staining and mNeonGreen fluorescence across pixels pertaining to each cell. GFP specking was quantified by local background subtraction (see below), high-pass filtering of fluorescent images, eliminating negative valued pixels, and calculating the mean fluorescence across pixels pertaining to each cell before and after high-pass filtering. Cells with low mNeonGreen or GFP expression were excluded from downstream analysis. For local background subtraction, images were down-sampled 8×8-fold. For each pixel in the full-resolution image, the local minimum across the 9×9 closest pixels in the down-sampled image was subtracted, after which negative values were eliminated.

### Analysis of HeLa genome-scale NIS-Seq perturbation screening data

Pairs of live phenotyping and NIS-Seq nuclear images acquired at corresponding stage positions were fine-mapped using FFT-accelerated cross-correlation. Phenotyping nuclei were assigned to the closest nucleus in shifted NIS-Seq data by centers of gravity, with a maximum movement distance of 11.1 μm and with the nuclear area matching within a twofold margin. Any ambiguously mapping nuclei were excluded from further analysis. Cells mapping to the same perturbed gene were aggregated by calculating the mean phenotype (e.g., the mean of Pearson correlations between nuclear and mNeonGreen signal in individual cells). Genes perturbed in less than 15 cells were excluded. Mean phenotypes from two independent replicate screens were visualized as scatter plots. Genes that displayed an altered phenotype in both independent screens were validated by individual lentiviral transductions.

### Analysis of THP1 genome-scale NIS-Seq perturbation screening data

THP1 phenotypes were imaged using a 10x objective with Z-stacking to better cover small ASC specks across the cellular cytosol. Furthermore, live nuclear imaging turned out to be affected by the stimulation of cells. Therefore, the assignment of phenotype and NIS-Seq images described above for HeLa cells was modified: Instead of using live nuclear images, cell outlines derived using CellPose from Z-aggregated membrane staining were shrunk by 5 pixels to predict the location of the nuclei. Potential pairs of phenotyping and NIS-Seq imaging fields-of-view were identified based on microscope stage positions. Images were scaled according to the relative magnification used, and coarsely mapped at 8×8-downsampled resolution using FFT-accelerated cross-correlation. Best-correlated pairs of fields-of-view were fine-mapped at 2×2-downsampled resolution using FFT-accelerated cross-correlation. All subsequent analysis steps were performed as described above for HeLa cells.

### Generation of single gene perturbation cell lines for validation assays

sgRNAs were selected from the Toronto human knockout pooled library (TKOv3), and ordered as DNA oligonucleotides (IDT, Supplementary Table 1). Oligos were phosphorylated and annealed at equimolar ratio with T4 PNK (Thermo) in 1x T4 DNA Ligase Buffer (Thermo) for 30 minutes at 37 °C, 5 minutes at 95 °C, and ramping to room temperature. Diluted annealed oligos were inserted into the CROPseq-iT7 lentiviral backbone using Golden Gate cloning. Purified and sequence-verified plasmids were used to produce lentivirus in HEK 293T cells. Transduced cells were selected with puromycin for 3-5 days and screened for the phenotype of interest.

## Data availability

Raw imaging data and example data will be available upon request.

## Code availability

All analysis tools are available as open-source web applications at *www.jsb-lab.bio/opticalscreening/*

